# Causality Analysis of Protein Corona Composition: Phosphatidylcholine-Enhances Plasma Proteome Profiling by Proteomics

**DOI:** 10.1101/2024.09.10.612356

**Authors:** Arshia Rafieioskouei, Kenneth Rogale, Amir Ata Saei, Morteza Mahmoudi, Borzoo Bonakdarpour

## Abstract

The study of the protein corona, the immediate and evolving biomolecular coating that forms on the surface of nanoparticles when exposed to a biological environment, is a crucial area in nanomedicine. This phenomenon significantly influences the behavior, functionality, and biological interactions of nanoparticles with biosystems. Until now, conclusions regarding the role of the protein corona in specific biological applications have been based on establishing *correlation* rather than *causation*. By understanding causality, researchers can predict how changes in nanoparticle properties or biological conditions will affect protein corona composition, in turn affecting the nanoparticle interactions with the biosystems and their applications. This predictive capability is essential for designing nanoparticles with specific characteristics tailored for therapeutic and diagnostic nanomedicine applications. Here, we explore the concept of “*actual causality*” (by Halpern and Pearl) to mathematically prove how various small molecules, including metabolites, lipids, vitamins, and nutrients, spiked into plasma can induce diverse protein corona patterns on identical nanoparticles. This approach significantly enhances the depth of plasma proteome profiling. Our findings reveal that among the various spiked small molecules, phosphatidylcholine was the actual cause of the observed increase in the proteomic depth of the plasma sample. By considering the concept of causality in the field of protein corona, the nanomedicine community can substantially improve their ability to design safer and more efficient nanoparticles for both diagnostic and therapeutic purposes.

## Introduction

Protein corona is a layer of evolving biomolecules, primarily proteins, that forms on the surface of nanoparticles when they interact with biological fluids, including human plasma^1,2^ This corona imparts a new biological identity to the nanoparticles, which becomes the point of contact with biosystems, including cells. The type, amount, and conformation of proteins on the nanoparticle surface determine how biosystems perceive and respond to the nanoparticles.^3,4^ Therefore, protein corona plays a critical role in dictating nanoparticle interactions with biosystems. Furthermore, the protein corona can significantly reduce the complexity of plasma proteins, thereby improving the detection capacity for low-abundance proteins.^5,6^ This is due to the protein corona’s unique ability to deplete highly abundant plasma proteins and enrich low-abundance ones on the nanoparticle surface.

While protein corona can reduce plasma complexity, the number of plasma proteins attached to nanoparticles is less than 10% of the total plasma proteins.^2^ To improve the number of identified proteins using the protein corona, several strategies have been developed, including the use of a “protein corona sensor array” with nanoparticles of distinct physicochemical properties.^5,6^ In this approach, various nanoparticles adsorb diverse set of proteins, which leads to enhanced proteome coverage. A recent strategy we developed involves spiking small molecules into plasma before incubation with nanoparticles.^7^ Small molecules such as nutrients, vitamins, lipids, and metabolites (namely, glucose, triglyceride, diglycerol, phosphatidylcholine (PtdChos), phosphatidylethanolamine (PE), L- α-phosphatidylinositol (PtdIns), inosine 5’-monophosphate (IMP), and B complex) can interact with plasma proteins, hindering their attachment to nanoparticle surface or altering their binding sites with nanoparticles and thus changing the protein corona composition.^7,8^

By applying this strategy, we observed a significant increase in the number of proteins attached to nanoparticles.^7^ Specifically, using protein coronas of polystyrene nanoparticles exposed to plasma treated with various small molecules (e.g., glucose, triglyceride, diglycerol, phosphatidylcholine, phosphatidylethanolamine, L - α-phosphatidylinositol, inosine 5-monophosphate, and B complex vitamins together with their combinations titled “sauce 1 and sauce 2”) at three different concentrations (i.e., 10, 100, and 1000µg/*ml*), we detected 1,793 proteins, marking an 8.25-fold increase compared to plasma alone (218 proteins) and a 2.63-fold increase relative to the untreated protein corona (681 proteins).^7^

Using mass spectrometry data from our recent study, here, we investigate the cause of the increased number of proteins using the rigorous mathematical concept of *actual causality*,^9^ rather than relying on statistical correlations. Almost all interpretations in the literature regarding the protein corona rely on correlation analysis (see **Fig. 1** for details). Our findings open up a new area in protein corona research where actual causality of compositional changes can be defined. This causality-based approach can enable the nanomedicine community to better predict the composition of protein corona when the physicochemical properties of nanoparticles change. For instance, this strategy can provide high-throughput opportunities for predicting the biological identity of various nanoparticle formulations designed for safe and effective genomic medicine delivery systems, where *in vivo* evaluation and comparison are challenging due to the expense and time required for large-scale animal studies.^10–14^ Understanding the actual causality of the effects of nanoparticles’ essential components (i.e., composition, physicochemical properties, and surface coatings) can help predict the biological fate and pharmacokinetics of nanomedicines, facilitating high-throughput screening and precise analysis of their biomolecular corona.^14^

**Figure 1.**
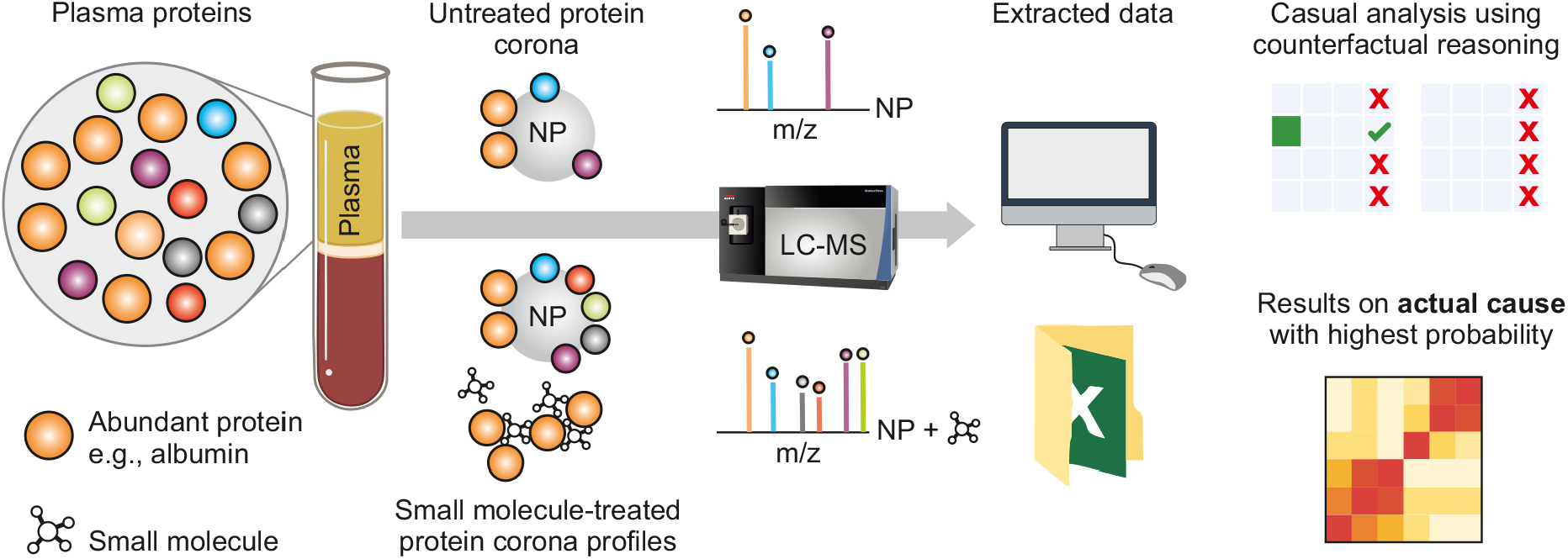
Scheme illustrating the overview of our work. The addition of small molecules to plasma can alter the interaction sites of proteins with nanoparticles, creating various corona profiles. Analyzing theses treated coronas with mass spectrometry technique significantly increases the number of quantified proteins when compared to untreated coronas or plasma. Actual causality analysis of the protein corona composition determines which small molecules are primarily responsible for enhancing the number of quantified plasma proteins in the protein corona.

## Actual Causality

Causality is a way of describing the logical dependence of events on one another. While there are multiple interpretations of the concept of causality, we focus here on the definition of *actual causality* as articulated by Halpern and Pearl. ^9^ This definition largely deals with what Halpern and Pearl refer to as token causality, which is causality regarding a specific occurrence of an event in a set of circumstances, rather than more generalized statements based on multiple observations.

Actual causality is based on *counterfactual reasoning*, which involves considering not only what actually occurred but also what might have happened under different circumstances. This type of reasoning, often encapsulated in questions such as “What might have happened if…?”, enhances our ability to imagine alternate scenarios (counterfactual worlds) as opposed to our current reality, which is referred to as the *actual world*. By imagining different outcomes based on varied actions or events, we build our capacity to understand and analyze the implications of those hypothetical changes.

Actual causality provides a mathematical framework to rigorously establish the causal effect of events in a system.

In many studies, including protein corona reports,^2,15–18^ statistical methods such as artificial intelligence (AI) and machine learning are being used to analyze experimental findings, often focusing on the correlation between events. This leads to the concept of correlation versus causation. Causal analysis seeks to identify the cause of an event, while correlation shows a statistical relationship. For example, during summer, both shark attacks and ice cream sales increase. While there is a correlation, there is no causation; the actual cause is the summer season itself, which drives both activities. One might mistakenly think that eating ice cream causes shark attacks, but the actual cause of both is the higher summer temperatures, which lead to both increased ice cream consumption and more people swimming in the ocean.

### Definition 1

*A basic causal model denoted as M, consists of variables and functions that map values to these variables. The notation* X *←* x *indicates that the variable* X *is assigned the value* x. *Likewise*, 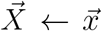 *denotes a vector of values* 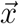 *to be assigned to a vector of variables* 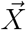

### Example 1

*Considering the causal model derived from the datasets obtained from experiments conducted in the report*,^*7*^ *we denote this causal model as M. In M, we have four features: Protein ID, Small Molecule, Concentration of Molecule, and log2FC (Log2 fold change). The variables in our causal model correspond to these features, and the function maps the domain (values obtained from samples) to these variables. For example, the variable* Concentration of Molecule *can take on three values:* 10, 100, *and* 1000µg/*ml. When we write* Concentration of Molecule *←* 10, *we are assigning the value 10 to the variable* Concentration of Molecule.

### Definition 2

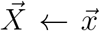 *is an* actual cause *of effect* φ *in causal setting M, if the following three conditions hold:*

- ***AC1***. *Both* 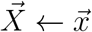 *and* φ *happen*.
- ***AC2***. *In the counterfactual world where we intervene on* X (X *←* x^*′*^*) in the actual world, the effect will not occur under the same contingency (the same setting as in AC1)*.
- ***AC3***. *The vector assignment* 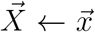 *is minimal; no subset of the value assignments in* 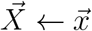 *satisfies AC1 and AC2*.

### Example 2

*In our running example (****Example 1****), the effect is measured by Log2 Fold Change (Log2FC). For highly abundant proteins, the threshold is set at Log2FC ⩾* 0.1, *whereas for other proteins, it is set at Log2FC ⩽* 0.1. *For instance, in our dataset, if a sample is listed among the highly abundant proteins and the Log2FC value is 2, the effect* φ *happens. Conversely, if a sample is not listed among the highly abundant proteins and the Log2FC value is 2, the effect does not happen*.

In this study, we are considering the *probabilistic* actual causation, which is the extension of Halpern and Pearl’s definition from Fenton-Glynn. ^19^ Fenton-Glynn extended Halpern and Pearl’s definition into a probabilistic version by adhering to the probability-raising principle, which is a traditional approach in the study of probabilistic causation. The core idea of this principle is that a cause should increase the probability of its effect. In this definition, we are changing **AC2** to a probabilistic version, called **PC2**. In this new definition, we state that the probability of the actual world occurring is greater than that of the counterfactual world. Specifically, the probability of the actual world occurring (where cause X *←* x and effect φ happen together) is greater than the probability of the counterfactual world occurring (where we intervene on X *←* x^*′*^ and effect φ happens) under the same conditions.

## Algorithmic Search for Actual Causality

Given a dataset of variables along with their values, we begin by searching through samples where the effect occurs, ensuring that **AC1** is satisfied. Within these samples, we calculate the probability of the effect occurring in both the actual world and a counterfactual world (where the feature’s valuation is altered). If the probability of the effect occurring in the actual world is higher than in the counterfactual world, the feature’s valuation satisfies **PC2**. In this study, we are seeking a singleton cause (a cause represented by a single variable), which means it is already in its minimal form and satisfies **AC3**. Consequently, we identify the actual cause for this subproblem. We have implemented this algorithm in the Python programming language.^20^

## Results and Discussion

Liquid chromatography mass spectrometry was used to analyze the protein corona compositions of various protein coronas formed after interactions between polystyrene nanoparticles and both untreated and small molecule-treated plasmas (respectively); full experimental details and data analysis is available in our recent report.^7^ Small molecules enhanced the detection of proteins by reducing the attachment of highly abundant plasma proteins to the nanoparticle surface while simultaneously increasing the attachment of low-abundance proteins. Cumulatively, we detected 1793 proteins with the array of small molecules, compared to 681 proteins in untreated proteins.^7^ As each of the small molecule could increase the numbers of detected proteins in protein corona, we were curious to identify the main cause for quantifying such a high number of the proteins. From the experimental outcome, we could only observe that PtdChos exerted a substantial increase in protein detection at the concentration of 1000µg/*ml*. To apply this concept to causality, we calculated the abundance ratio of each protein in the corona of untreated plasma to that of treated plasma and presented their log2FC values (see the **Supplementary Data 1** for details; explanations on the treatment of the missing values for some small molecules are provided in the original research^7^) for our causality analysis, alongside three other features: Protein ID, Small Molecule, and Concentration of Molecule. A list of descriptions for the variables can be found in **Table. 2**. In the causal model, these features serve as variables, with the effect measured by Log2FC (Log2 fold change), which has a threshold of 0.1 or more for highly abundant proteins, (as listed in **Table. 1**), and 0.1 (or less) for the rest of the identified proteins in protein corona, highlighting the need for depletion of highly abundant proteins and enrichment of the low abundance proteins.

**Table 1.**
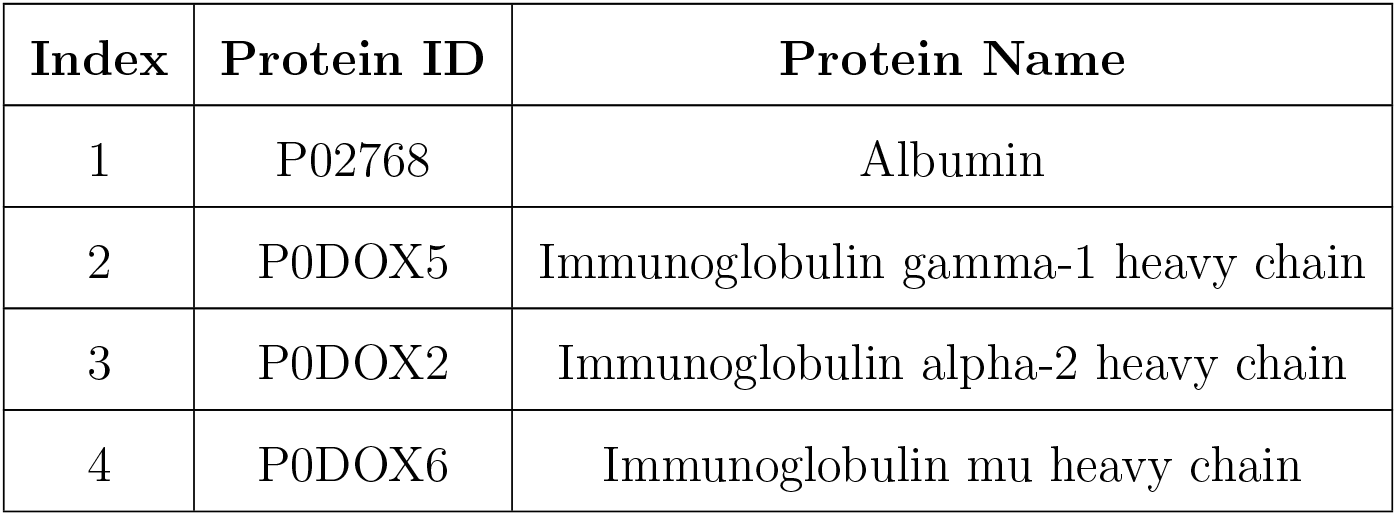
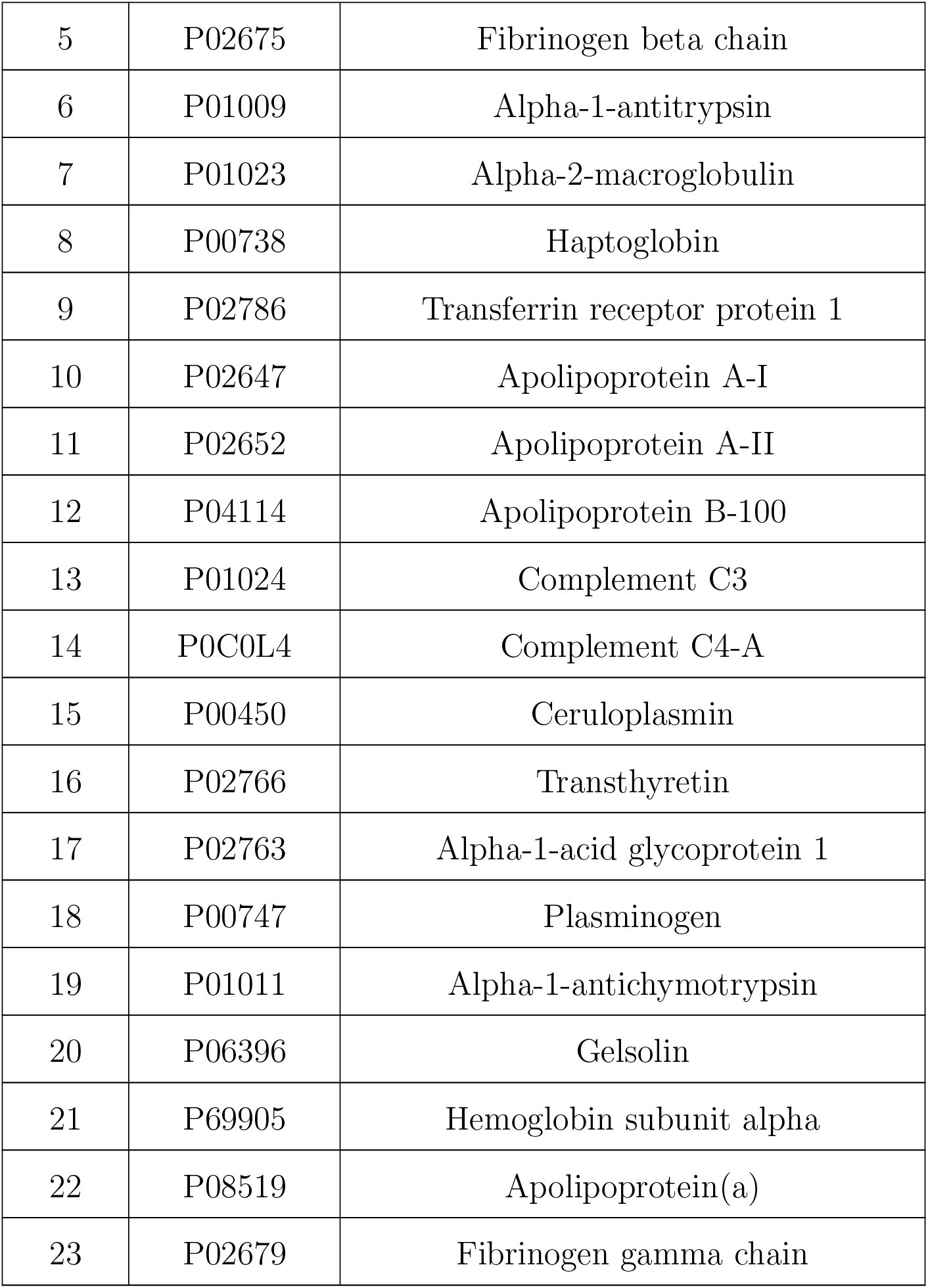
List of highly abundance proteins.

**Table 2.**
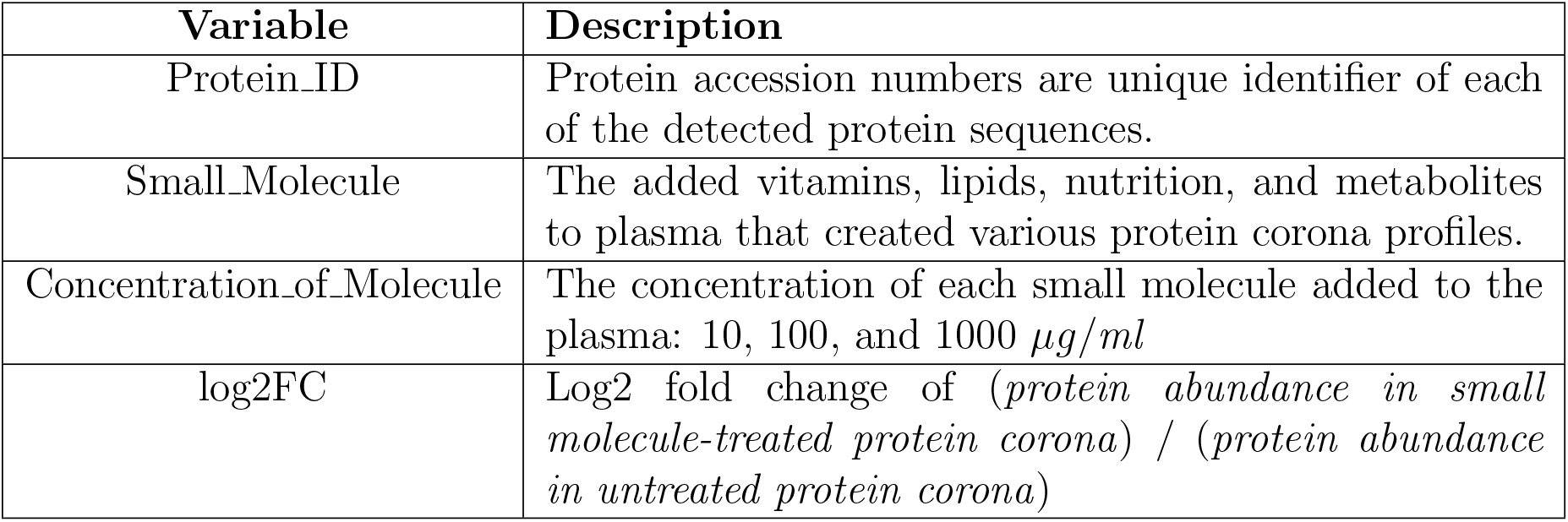
Description of Variables Causal Model.

| Variable | Description |
| --- | --- |
| Protein_ID | Protein accession numbers are unique identifier of each of the detected protein sequences. |
| Small_Molecule | The added vitamins, lipids, nutrition, and metabolites to plasma that created various protein corona profiles. |
| Concentration_of_Molecule | The concentration of each small molecule added to the plasma: 10, 100, and 1000 $\mu\text{g/ml}$ |
| log2FC | Log2 fold change of (*protein abundance in small molecule-treated protein corona*) / (*protein abundance in untreated protein corona*) |

Our objective in this approach was to identify the actual causes by examining various threshold combinations for highly abundant proteins. We divide the spectrum into two segments: thresholds between the minimum value of Log2FC and 0.1 for highly abundant proteins, and thresholds between 0.1 and the maximum value of Log2FC for the remaining proteins. At each step, we establish two thresholds—one for highly abundant proteins and one for the remaining ones. The effects were then determined based on these thresholds, allowing us to systematically investigate the impact of different threshold combinations on the cause that we found.

In **Fig. 2**, the procedure for finding causation using counterfactual reasoning is illustrated. The left table represents the actual world, where we search for samples (values assigned to variables) in which the effect has occurred. The right table represents the counterfactual world, where we search for samples that differ only in the variable of interest, while keeping the other variables the same. We then calculate the probability of the effect occurring in the actual world compared to the counterfactual world. If the probability in the actual world is greater than in the counterfactual world, the cause can be identified as the value assigned to the variable being examined. For example, in **Fig. 2**, the cause is *Small Molecule ← PtdChos*.

**Figure 2.**
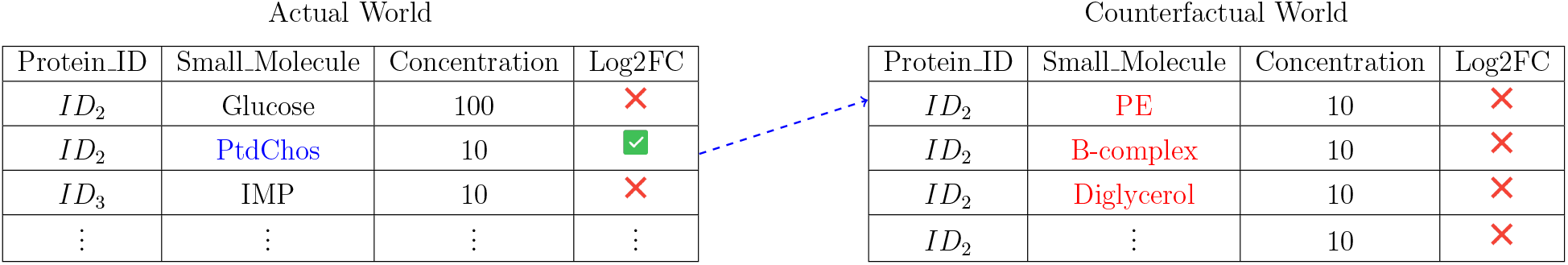
Procedure of counterfactual reasoning. The left panel illustrates the actual world and the right panel depicts the counterfactual world.

In **Fig. 3** we determine the actual cause with the highest, second highest, and third highest probabilities by applying our mathematical approach. This method employs a low threshold for highly abundant proteins and a high threshold for the low abundance proteins. In each cell shown in **Fig. 3**, the causes with the highest, second highest, and third highest probabilities are identified, respectively (**Fig. 3a, Fig. 3b**, and **Fig. 3c**). Our results revealed that more than 50% of the causes with the highest probability are related to the spiking of `PtdChos`. Further analysis revealed that in over 65% of the settings with different thresholds, `PtdChos` is among the top three causes with the highest probability. This outcome is particularly intriguing, as the experimental correlation data only indicated the critical role of `PtdChos` at the highest concentration used (i.e., 1000µg/*ml*). In contrast, the causality results demonstrated the critical role of `PtdChos` independent of its concentration.

**Figure 3.**
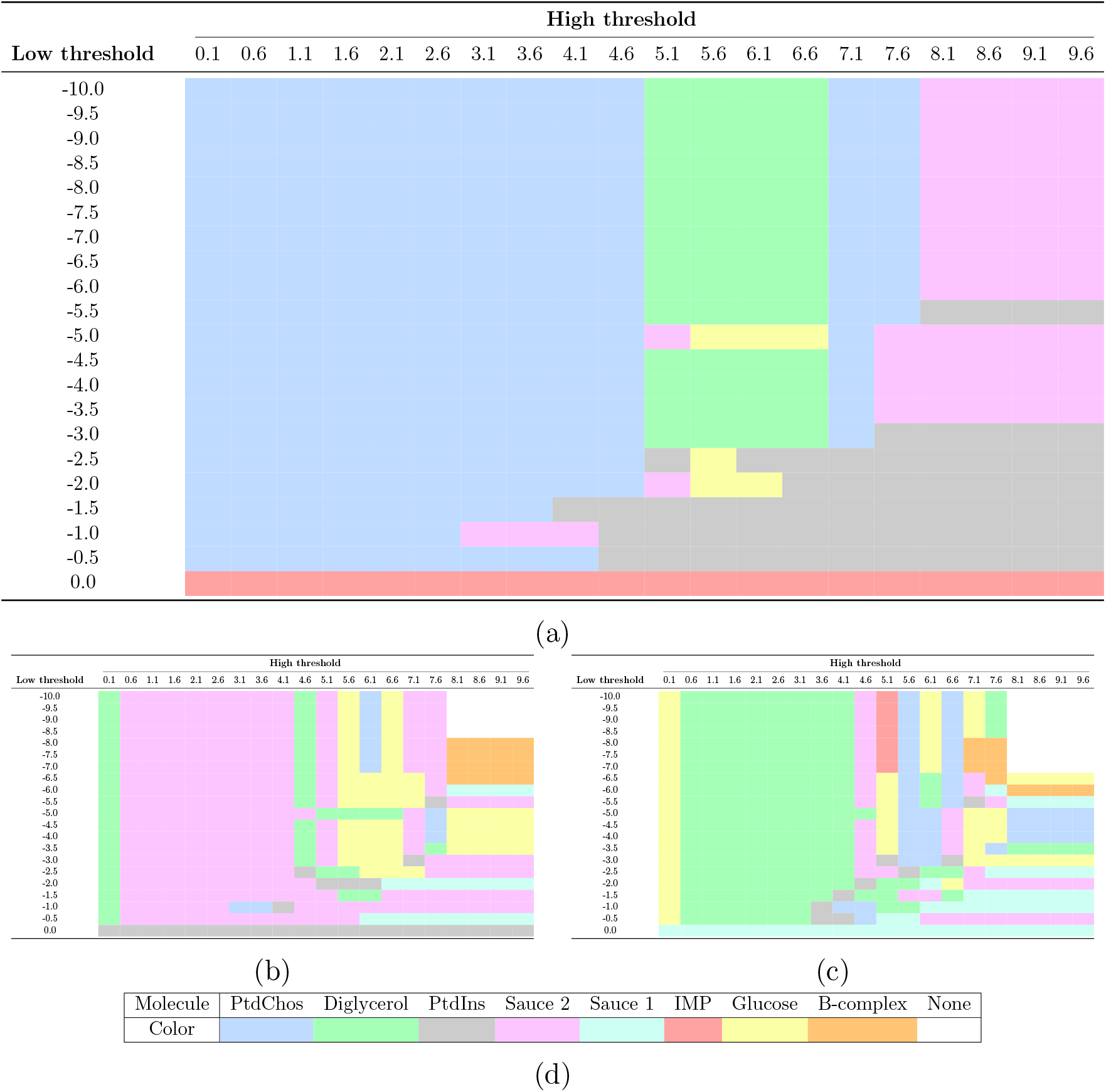
Actual cause with the (a) highest, (b) second highest, (c) and third highest probabilities in different threshold settings. Details about the colors of each cell can also be found in (d). The employed molecules in this study are: `glucose`, `triglyceride`, `diglycerol`, phosphatidylcholine (`PtdChos`), phosphatidylethanolamine (`PE`), L-α-phosphatidylinositol (`PtdIns`), inosine 5-monophosphate (`IMP`), and vitamin `B-complex`. `Sauce 1` and `Sauce 2` are combinations of four small molecules each. Molecular `Sauce 1` contains `glucose, triglyceride, diglycerol`, and `PtdChos`, while Molecular `Sauce 2` consists of `PE`, `PtdIns`, `IMP`, and vitamin `B-complex`. Additionally, ‘‘`None`’’ indicates that no cause with any probability was found for that cell with the given threshold setting.

In summary, we have introduced the concept of actual causality and demonstrated its critical importance over statistical correlation in mechanistically understanding the causes of compositional changes in the protein corona. Specifically, we examined the intricate dynamics of the protein corona, focusing on the role of small molecule-spiked plasma in enhancing nanoparticle efficacy in proteomics. By employing a causality-based approach, we moved beyond traditional correlation methods to rigorously define the actual causes of these compositional changes. Our findings revealed the significant impact of `PtdChos` in the protein corona, highlighting its critical role independent of concentration levels. This actual causality analysis provides a more robust understanding of how specific small molecules influence nanoparticle interactions with biological systems.In addition, it provides a unique oppurtunity to look into the derivatives of small molecules of actual cause to even more improve the deep proteom profiling of plasma. This causality-based perspective in the field of protein corona offers a high-throughput predictive capability, enabling more precise design of with predictable protein coronas nanoparticles tailored for therapeutic and diagnostic purposes. As we continue to refine our understanding of the protein corona, the insights gained from this study pave the way for safer, more effective nanomedicine applications, ultimately contributing to the advancement of personalized medicine.

## Supporting Information

The Supporting Information is available free of charge at link ^1^. Supplementary Data 1 and Supplementary Data 2 provide details of the mass spectrometry outcomes and the statistical analysis of the causality of top three small molecules in each of the predetermined thresholds (respectively).

## Data and code availability

The developed code and related data are available through the following link^2^

## Footnotes

1 *https://???*

2 *https://github.com/arshiarafiei/Causality_Analysis_of_Protein_Corona*

